# From 1D sequence to 3D chromatin dynamics and cellular functions: a phase separation perspective

**DOI:** 10.1101/255174

**Authors:** Sirui Liu, Ling Zhang, Hui Quan, Hao Tian, Luming Meng, Lijiang Yang, Huajie Feng, Yi Qin Gao

## Abstract

The high-order chromatin structure plays a non-negligible role in gene regulation. However, the mechanism for the formation of different chromatin structures in different cells and the sequence dependence of this process remain to be elucidated. As the nucleotide distributions in human and mouse genomes are highly uneven, we identified CGI forest and prairie genomic domains based on CGI density, which better segregates genomic elements along the genome than GC content. The genome is then divided into two sequentially, epigenetically, and transcriptionally distinct regions. These two types of megabase-sized domains spatially segregate, but to a different extent in different cell types. Overall, the forests and prairies gradually segregate from each other in development, differentiation, and senescence. The multi-scale forest-prairie spatial intermingling is cell-type specific and increases in differentiation, thus helps define the cell identity. We propose that the phase separation of the 1D mosaic sequence in space, serving as a potential driving force, together with cell type specific epigenetic marks and transcription factors, shapes the chromatin structure in different cell types and renders them distinct genomic properties. The mosaicity of the genome manifested in terms of alternative forests and prairies of a species could be related to its biological processes such as differentiation, aging and body temperature control.

## Introduction

Eukaryotic chromatins possess highly complex structures which are of great biological importance. The heterochromatin compaction and the cell-or tissue-specific activation together shape the chromatin. On one hand, the folding of chromosomes must be robust in order to protect the genetic material. On the other hand, flexibility is needed to allow different DNA sequences to be accessed in response to different signals. Using Hi-C and ChIA-PET techniques, recent studies have shown that the 3D chromatin structure is important for gene regulation^1, 2^. Our comprehension of genome architecture has since advanced rapidly in recent years, resulting in identification of structural domains at different scales (e.g. loops^3, TADs4-6^, types^7^, and compartments^1^) and a better understanding of their roles in gene regulation. Much progress has been made in the chromatin structural study of different cell types^8, 9^ and different cellular processes like early embryonic development, cell differentiation, and cell senescence^10-15^.

Multiple factors contribute to the chromatin structure formation and functioning of organisms. For example, HP1 and polycomb proteins bind to H3K9me3 and H3K27me3 repressive histone marks, respectively, to form constitutive and facultative heterochromatins. CTCF, previously recognized as a transcriptional insulator that blocks enhancer-promoter interactions^16, 17^, is reported to be enriched at TAD boundaries and its knockdown leads to an increase in inter-TA D interactions^4, 18^. Loss of cohesin protein which is recruited by CTCF also leads to interaction increase between neighboring TADs, despite that the impact seems less than that of CTCF^18, 19^. In mitosis, “mitotic bookmarking” transcription factors have been suggested to play a role in chromatin structure re-establishment^20^. These factors along with epigenetic modifications shape the chromatin structure of different cell types via specific or non-specific binding to sequences.

Gene positioning and transcriptional activity represent major determinants of the microscopic chromatin structure that self-organizes in a rather predictable way. However, there is much to learn about the primary DNA sequence as the “footprint” of DNA structure and packaging. The DNA coding sequence only accounts for less than 5% of the mammalian genome, and the role of the rest of the genome is largely unknown. Though their specific function is largely under debate, noncoding DNAs are increasingly believed to play an architectural role in the formation of complex eukaryotic chromatin. Efforts have been paid to investigate the relationship between the mosaic, multi-scale genomic sequences and the spatial structure of chromatin. Grosberg et al associated the long-range correlations of the DNA primary sequences with their 3D structures^21^.

In particular, the genomes of warm-blooded vertebrates are known to display alternations between AT-rich and GC-rich homogeneous genome regions called isochores, which have distinct biological properties including gene density and replication timing^22, 23^, and were reported to associate with TADs and Lamina Associated Domains (LADs)^24^. Besides the isochores, CpG dinucleotides also tend to aggregate to form CpG islands (CGIs). They usually locate at the promoter regions of genes and play important roles in gene expression regulation. CGIs at the promoter regions of genes are involved in gene regulation via hypermethylation and binding of transcription factors and regulatory proteins such as polycomb complex^25^.

In this study, we analyze the uneven distribution of CGIs along the genome, and investigate how this mosaicity of the DNA sequence affects the packaging and thus functioning of the genome under different cellular conditions. We found that the human and mouse genomes can be divided into large (megabase scale) alternative domains of high and low CGI densities, named forests (F) and prairies (P), respectively. This division partitions the genome into two types of regions that are genetically, epigenetically, and transcriptionally distinctly different, and outperforms isochores in the segregation of these properties. More importantly, interactions and package of forests and prairies in the 3D space show consistent changes during the process of early embryonic development, cell differentiation and senescence. The spatial segregation of prairies from forest indicates a phase separation mechanism in chromatin structure formation and re-modelling, and the lineage specific interaction between the two types of DNA domains in cellular processes provides a new view on how cellular functions are achieved through the control of chromatin 3D structures. Lastly, we discuss the possible physical reasons behind the domain segregation, how the F-P alternative mosaic genome might affect the biological function of related species, and how the sequence difference between different species might be related to chromatin function realization.

## Materials and Methods

### Definition of CGI forest and prairie

The forests are defined as DNA domains with densely distributed CGIs and prairies low CGI densities. We identified CGI forests and prairies based on neighboring CGI distances along the genome. We first defined a critical neighboring CGI distance for the genome under study, longer than which the neighboring CGIs are more enriched than by random chance. It is noted here that the critical distances vary with the chromosomes, reflecting their CGI densities and clustering patterns. A CGI forest was then defined as a continuous DNA region longer than the critical length, inside which all neighboring CGI distances are shorter than the critical length. Prairies were defined as the complement to forests in each chromosome excluding the longest chromosomal gap.

### An alternative forest and prairie definition

To evaluate the robustness of the F-P definition over CGI identification, we defined CGI in an alternative way and examined the overlap between forests identified according to the canonical CGIs obtained from UCSC (http://genome.ucsc.edu/cgi-bin/hgTables) and the newly defined CGIs. The new definition of CGI was given based on the CpG density of each 200bp window using a sliding window approach. A window was defined as a CGI if its CpG density is greater than 0.075. The adjacent CGIs were merged into a larger one. The alternative CGI F-P definition was based on the neighboring distances between these new CGIs using the same method described above.

### Enrichment of histone marks, DNase I hypersensitive sites (DHS) and Ogt protein binding sites (OBS) in forests and prairies

The data for histone modification, DHSs and OBS were obtained as densities of the corresponding signals (raw signal for OBS and fold-change for the rest). The enrichment value for DHS, OBS and each histone mark of individual forest or prairie is defined as the average signal

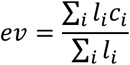

where *c*_*i*_ is the signal of the *i*th fragment located in the domain, and *l*_*i*_; the length of the *i*th fragment. We analyzed the enrichment levels of 28 histone marks for IMR90 cell line and 5 core histone marks (H3K4me1, H3K4me3, H3K9me3, H3K27me3 and H3K36me3) in various samples, including those from human brain tissues, blood cells, normal somatic tissues and cell lines.

### Enrichment of transcription factors in forests and prairies

The transcription factor binding sites (TFBS) for different transcription factors (RNA polymerase II, Cebpb and Rad21) were downloaded in the narrowPeak format from UCSC genome browser. The enrichment of each TF in each forest and prairie was evaluated by peak density (*pd*) defined as the ratio of the peak number in the domain to the domain length

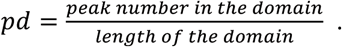

### F-P difference for epigenetic properties

To quantify the difference between forests and prairies for each epigenetic feature, including methylation level, histone marks, DHS, OBS and TFBS enrichment, we defined the epigenetic signal difference of each domain as

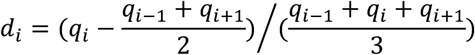

where *q*_*i*_, *q*_*i-1*_ and *q*_*i+1*_ are the epigenetic quantities for the *i*th domain and its two flanking domains.

### Enrichment ratio for histone marks

The enrichment ratio of histone marks was defined for forests and prairies, respectively, as the ratio between the number of domains with positive enrichment differences (*d*_*i*_) and the total domain
number

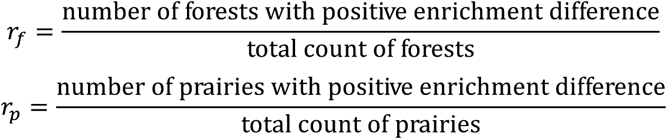

A larger *r*_*ƒ*_ (*r*_*p*_) indicates that more forests (prairies) enrich the corresponding histone mark.

### Type definition

In our previous work^7^, we defined chromatin type A and B structures according to their local Hi-C contact patterns with a Linear Discriminant Analysis (LDA) classifier based on PMD/non-PMD identification. We improved the method by both modifying the classifier and expanding the type A/B definition to the whole genome. For each sample, we identified typical types A and B based on the peak numbers of the local contact summation in 800-kb range for each 800-kb unit along the sequence. A neural network classifier was then trained on typical types A and B and each 800-kb unit was scored with a sliding window of 40 kb. Neighboring units classified into the same type were merged together. In this way, the structure type is identified purely according to their local structures. This method gives an average accuracy of 93% in 5-fold validation for IMR90 cell line. All Hi-C data in this work were normalized by ICE normalization in a 40-kb scale using the iced python package^26^.

### Compartment identification

We defined compartments A and B following Lieberman-Aiden’s approach^1^ with slight modifications. We first calculated the intra-chromosomal observed/expected matrix and performed Eigen decomposition on the correlation matrix of the corresponding Z-score matrix. To decide which eigenvector to use, we defined a parameter

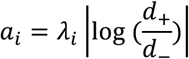

for the *i*th dimension, where *λ*_*i*_ is the *i*th largest eigenvalue, *d*_+_ is the gene density in regions with positive entries of the corresponding *i*th eigenvector, *d*_ is the gene density in regions with negative entries, and chose the eigenvector with the highest *a*_*i*_ among the first three dimensions.

### Forest index calculation

To quantify the local structural environment of forests and prairies, we defined a forest index *ƒ*_*i*_ for 40-kb bead *i* as

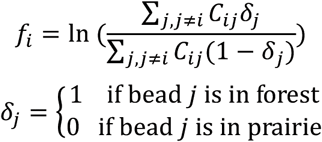

where *C*_*ij*_ is the normalized Hi-C contact probability between beads *i* and *j*. The self-contact was excluded in the calculation. For each 40-kb bead, more frequent interactions with forests than with prairies render a positive forest index. A higher absolute value of *ƒ*_*i*_ indicates a more uniform environment. As the contact probability decays in a power-law form along the genomic distance, local interactions naturally contribute more to the forest index than long-range interactions.

### Domain contact types and their proportion

Three contact ratios were calculated for forest and prairie domains, respectively, based on the domain contact matrix, whose entry *D*_*ij*_ represents the sum of contacts between domains *i* and *j*. The self-contact on the diagonal of the 40-kb resolution Hi-C matrix was subtracted before the calculation of domain contact matrix. The intra-domain contact ratio was calculated as

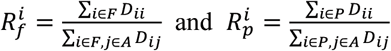

for forests and prairies, respectively, in which *F* is the collection for all forest domains, *P* is the collection for all prairie domains, and *A* is the union of sets *F* and *P*. The inter-domain contact ratio between the same domain types was calculated as

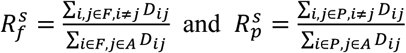

and the inter-domain contact between different types similarly as

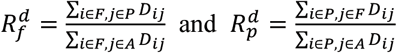

### Segregation ratio calculation

The segregation ratio *R*_*s*_ was defined as the ratio of inter-domain contacts between the same types and different types

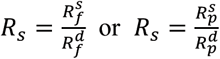

for forests and prairies, respectively.

## Results and discussion

### Forests and prairies are large genomic domains with distinctly different genetic features

Similar to CpG dinucleotides which linearly segregate to form CpG islands (CGIs), CGI distribution is also uneven along the DNA sequence (Figures 1A and S1). Here we define CGI forest (F) and prairie (P) domains based on neighboring CGI distances. A CGI forest is rich in CGIs, while a CGI prairie is where CGIs are sparse, as their names suggest (Table S1). Forests and prairies in human show similar megabase-scale average lengths and their length ratio varies by chromosome (Figure S2). In mouse, their average lengths are slightly longer, especially for prairies (Table S1). The identification of CGI forests and prairies is robust over CGI definition (Figure S3).

**Figure 1.**
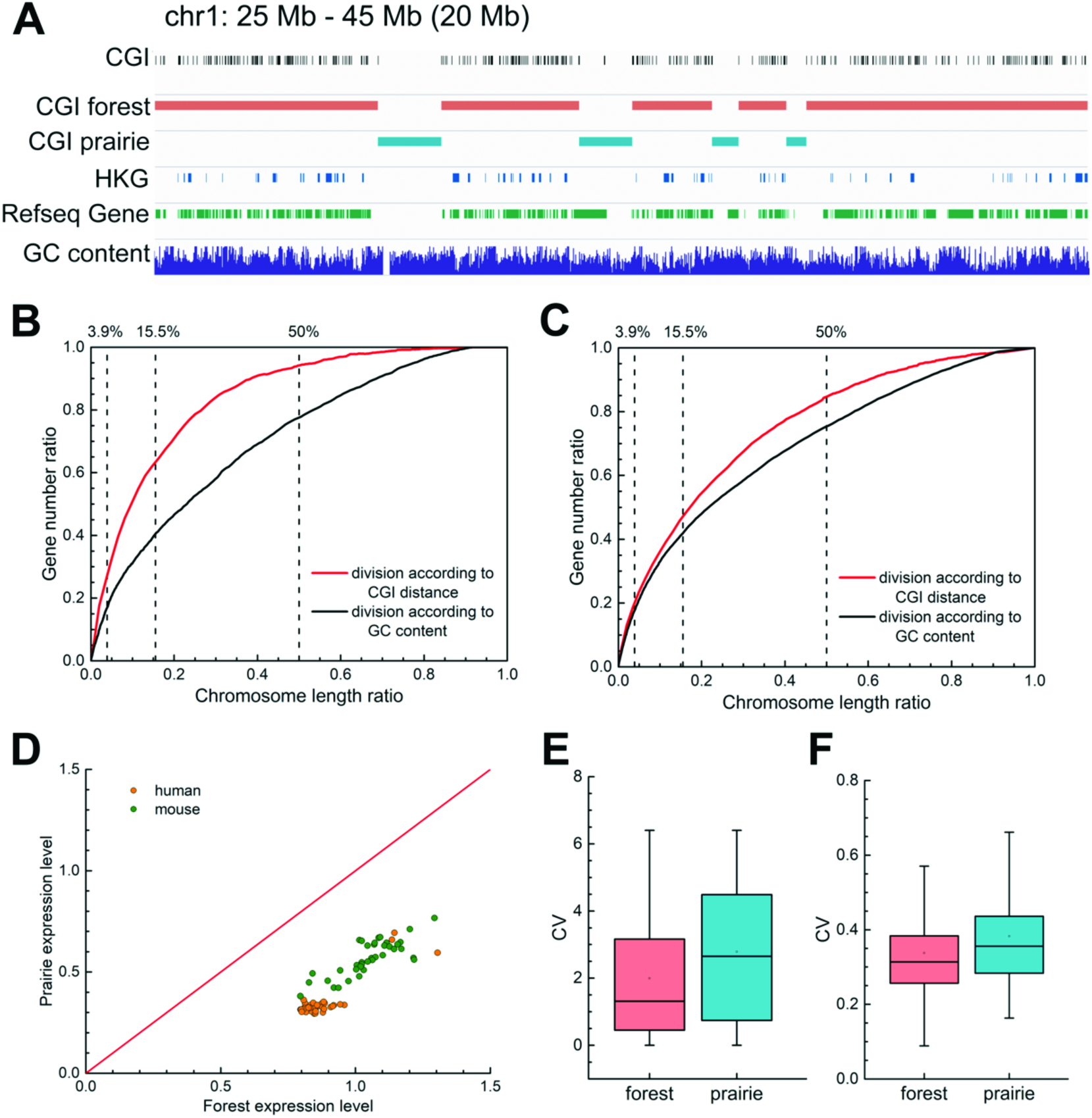
The genetic features of CGI forests and prairies. (A) IGV snapshot for a representative 20-Mb region on human chromosome 1 showing that CGI forests are where CGIs cluster, and are enriched in genes, especially housekeeping genes. (B) the characteristic curve of the division by CGI distance and by GC content in regard to housekeeping genes and (C) all genes. For GC content, each point on the curve shows the length ratio of regions with GC content above a threshold and the proportion of genes in these regions. For CGI distance, each point shows the length ratio of regions with neighboring CGI distances lower than a threshold, and the corresponding ratio of genes located inside. A higher AUC means that the feature is enriched at a shorter chromosome length, thus indicating a more effective feature enrichment strategy. (D)Mean gene logarithm expression levels in forests and prairies of different human and mouse samples. (E) The boxplot for CVs of expression levels in human samples for all genes and (F) for housekeeping genes.

CGI forests are enriched in genes, especially in housekeeping genes (HKGs). Despite their shorter total length than prairies, forests possess 91.3% of the HKGs and 78.5% of all genes, with an overall gene density 3.7 times higher than that in prairies (Table S1). Although 72% gene promoters are CGI promoters^27^, 63.3% genes with non-CGI promoters also reside in forests, indicating that gene enrichment in forests is not simply caused by the clustering of CGI promoters. The mouse genome is of similar properties (Table S1).

To assess to what extent the F-P division can dissect the genome by genomic features, we gave the feature enrichment characteristic curves of F-P division for HKGs and all genes, and compared them with the performance of GC content (Figures 1B and 1C). CGI distribution’s characteristic curve in regard to HKGs is significantly higher than that of GC content (Figure 1B and Table S2). Its area under the curve (AUC) is also noticeably higher (0.843 for CGI distance, and 0.709 for GC content). When all the genes are considered, the F-P classification still outperforms the GC content division in gene segregation (Figure 1C and Table S2), with higher AUC (0.754 vs 0.700) and higher gene ratio at the same length. Similar results are also obtained for the mouse genome (Figure S4). Therefore, gene-rich/poor regions segregate more distinctively according to CGI density than to GC contents. Genes in forests and prairies are distinct in biological functions. For example, Gene Ontology (GO) analysis using DAVID (https://david.ncifcrf.gov) ^28, 29^ shows that HKGs in prairies are specifically enriched in GO terms of DNA damage and repair, chromatin remodeling, p53 signaling, and cellular response to oxidative stress compared to those in forests (Figure S5).

The gene expression levels are affected by the genome landscape. Genes in forest are significantly more highly expressed but varies less across cell types than those in prairies (both with p-value < 10^−100^ by Welch’s unequal variance t-test for logarithm expression and the coefficients of variation (CV), Figures 1D and 1E). However, HKGs in forests and prairies are similar, both possessing higher average expression levels and varying notably less than all genes (Figures 1F and S6). In contrast, tissue-specific genes in both forest and prairie vary among cells significantly more than all genes (p-value < 10^−100^ by Welch’s unequal variance t-test). Tissue-specific genes in prairies also vary significantly more than those in forests (with CVs of 2.33 and 1.65, respectively). The higher variances for genes in prairies indicate that they are more extensively regulated than forests, thus may play an important role in cell differentiation, as are validated later.

Although CGI forests and prairies exist in vertebrates like human and mouse, the CpG density distribution can vary greatly among species. For example, the CVs of the CpG densities of invertebrate *Drosophila melanogaster*, plant *Arabidopsis thaliana*, single-celled organism *Schizosaccharomyces pombe*, and bacteria *Caulobacter crescentus*, are 0.146, 0.176, 0.096, 0.111, respectively, and are all significantly smaller than that of human (0.578) and mouse (0.463). These genomes have high and uniform CpG distributions, which can thus be considered as consisting of mainly forests, with little mosaicity (Figure S7).

### The epigenetic features of forests and prairies

Besides genetic features, many epigenetic features are also consistently different between forests and prairies, including DNA methylation, histone modifications, DNase I hypersensitive sites (DHSs), and transcription factor binding sites (TFBS) (Figure 2A). Interestingly, the discrepancy between forests and prairies varies significantly with cell type, showing a consistent change following cell differentiation and development.

**Figure 2.**
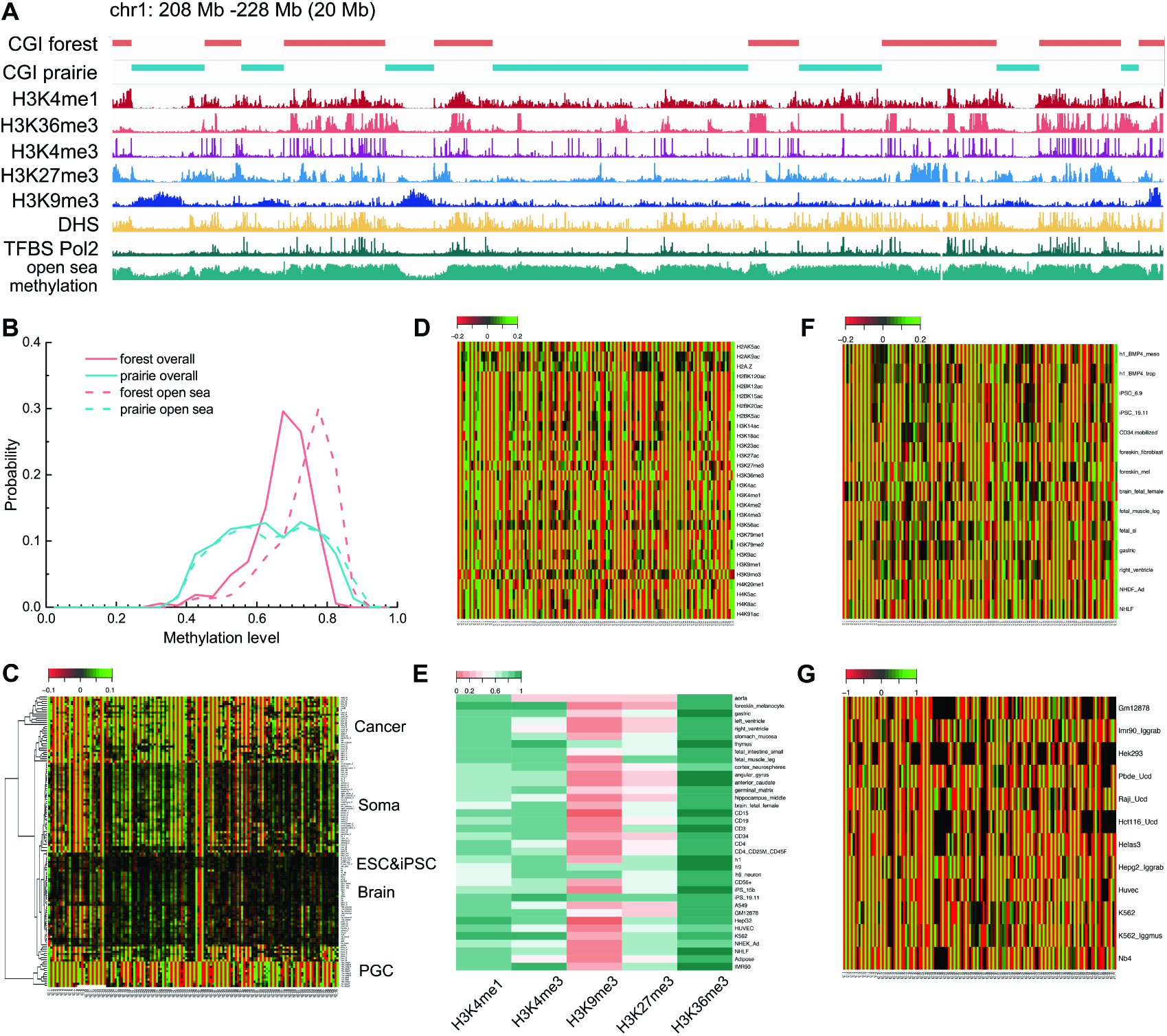
The epigenetic features in forests and prairies. (A)IGV snapshot for a representative 20-Mb region on human chromosome 1 showing the open sea DNA methylation level, distribution of histone marks, DHS and TFBS in forests and prairies of IMR90 cell line. (B) The probability distribution of the overall and open sea methylation level in forests and prairies for IMR90 cell line. (C) Heatmap of F-P methylation level difference in different cell types. Columns represent forest and prairie domains in the sequential order, and rows are samples hierarchically clustered according to their methylation difference patterns. (D) Heatmap of F-P enrichment difference of different histone marks for IMR90 cell line. (E) Enrichment ratio for the 5 core histone marks (H3K4me1, H3K4me3, H3K9me3, H3K27me3 and H3K36me3). (F) F-P DHS distribution difference in different samples. (G) F-P density difference of TFBS for RNA polymerase II (Pol2) in different samples. (Data given here are for human chromosome 1)

Since the methylation of CGIs usually associates with the specific regulation of CGI-promoter genes, their methylation level is actively controlled and normally remains low. In contrast, the methylation level of the open sea (defined as the genomic regions excluding CGIs, CGI shores and CGI shelves^30^) lacks specificity and better reflects the environmental chromatin state. We found that, in general, the overall methylation level of forests is lower than that in prairies, while the open sea methylation level shows an opposite trend in almost all samples examined except for the primordial germ cells (PGCs) which undergo demethylation (Figures 2B and S8)^31^.The open sea and overall methylation level change around forest-prairie boundaries is in good agreement with that of the whole domains (Figure S9). Forests and prairies also yield a clearer discrimination of human genome regarding CpG methylation than isochores (Figure S10).

We next calculated the open sea methylation level difference between neighboring domains (see method). The result given in Figure 2C clearly shows a negative-positive-alternating pattern, with the open sea methylation level of a forest almost always higher than that of its neighboring prairies. This pattern is conserved in most cell types except for PGCs, which have lower open sea methylation level in forests than that in prairies (Figures 2C and S8). The apparent methylation difference between forests and prairies indicates that they separate the genome into epigenetically distinct domains, and can be regarded as functional and structural units. Hierarchical clustering of the methylation differences across different cells demonstrates their cell-type specificity. The F-P methylation difference is small for brain cells, embryonic stem cells (ESCs), induced pluripotent stem cells (iPSCs) and gonadal somatic cells, large for cancer cells and immortalized cell lines, and intermediate for somatic cells. Notably, as shown in previous study^32^ as well as discussed later, the different methylation levels of forests and prairies also correlate to their differences in the spatial packing of the chromatin.

To further evaluate the chromatin states of forests and prairies, we investigated the F-P difference of histone marks^33^, whose enrichment difference also shows a positive-negative alternation at the domain level. Active histone marks such as H3K4me1, H3K4me3, and H3K36me3^34^ concentrate in forests, whereas repressive histone mark H3K9me3 aggregates in prairies (Figures 2D and 2E). The enrichment difference of histone marks in other cells shows a similar pattern, but with a significant cell-type dependence (Figures S11 and S12). In particular, the conventional repressive mark H3K27me3 shows a preference for forests or prairies in a cell-type specific way.

Moreover, the DNase I hypersensitive site (DHS, an open chromatin signal ^35^) are significantly more enriched in forests than in neighboring prairies for all cell types studied here, suggesting that forests adopt a more open conformation with higher chromatin accessibilities (Figure 2F). The transcription factor binding site (TFBS) density for RNA Polymerase II in forests is also higher than that in prairies, so are the densities for other TFBSs such as Cebpb (CCAAT enhancer binding protein) and Rad21 cohesin (Figures 2G and S13). The notion that the forests largely constitute the open chromatin is further validated by the significant enrichment of the O-linked N-acetylglucosamine (O-GlcNAc) transferase (Ogt) in forests (p-value<10^−82^ by Welch’s unequal variance t-test, Figure S13), which as a DNA binding protein modifier does not bind to DNA sequence directly^36^. Therefore, the distributions of DNA methylation, histone marks, DHSs and TFBSs in forests and prairies all show that forests and prairies segregate the chromatin into distinct epigenetic domains, and the open and active chromatin is formed mainly by forests rather than prairies.

### Forests and prairies have distinct structural properties

Next, we examined the structural properties of forests and prairies. Based on the Hi-C data, we found that forests and prairies form different 3D structures. First, F-P boundaries largely overlap with TA D boundaries. Isochores were also previously reported to overlap with TAD boundaries significantly better than random^24^. We found that although both F-P boundaries and isochore overlap with TAD boundaries (Figures 3A and 3B), the former have a much higher significance level (p-value<10^−24^ by chi-square test) than the latter (0.02). As TAD boundaries were previously reported to function as insulators and exhibit distinct properties at opposite sides^4^, their cooccurrence with F-P boundaries suggests the roles of the latter in segregating genomic and structural domains.

**Figure 3.**
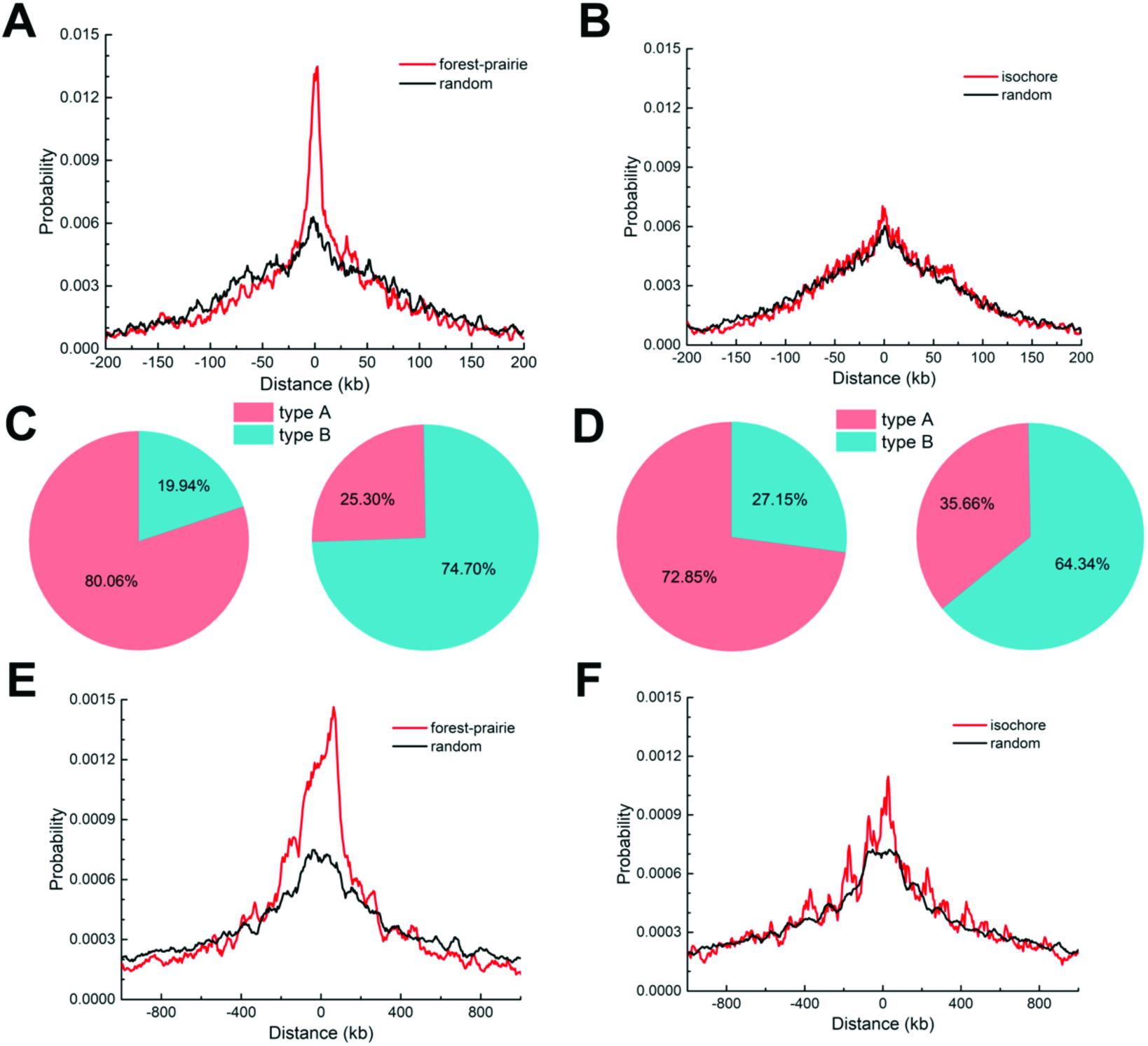
Structural properties of forests and prairies. (A) Distance distribution of F-P boundaries to TAD boundaries and (B) isochore boundaries to TAD boundaries. (C) The composition of type A and B in forest (left) and prairie (right), and (D) in high GC content regions (left) and low GC content regions (right). (E) Distance distribution of F-P boundaries to compartment boundaries and (F) isochore boundaries to compartment boundaries.

We then examined the local structures formed by individual forests and prairies. The type A and type B regions are regarded as chromatin secondary structures and mainly comprise compartments A and B, respectively^7^. We expanded the previous type A/B definition to the whole genome using a neural network classifier (see method), with types A and B consist of 46.7% and 45.2% of the whole chromosome, respectively. Despite their similar average lengths, type As have a significantly higher average TAD number, gene density, and expression level (p-value < 10^−43^, 10^−22^, and 10^−22^ by Welch’s unequal variance t-test), and are less categorized into the “unstructured regions”^3^ (Table S3). We next examined the structures of forests and prairies, and found that 80.06% of forests are type A, and 74.70% prairies type B. In contrast, lower proportions (72.85% and 64.34%) of high and low GC content regions are made up of type A and B, respectively (Figures 3C and 3D). These results show that CGI distribution (prairie and forest) is highly predictive for chromatin 3D structure formation.

Chromatin 3D structures are commonly partitioned into compartments. We defined compartments following Lieberman-Aiden et al’s procedure^1^ with slight modifications, and compared the compartments of different cell types with forest and prairies for both human and mouse samples. We found that on average 67.7% forests and 71.5% prairies of human, and 80.0% and 81.2% for mouse, lie in compartments A and B, respectively. The common compartments A and B for all the samples analyzed are also mainly composed of forests and prairies, respectively (SI). The forest-prairie boundaries significantly enrich at compartment boundaries for both human and mouse (p-value < 10^−37^ and < 10^−30^ by chi-square test, Figures 3E and S14). In contrast, the segregation of high (Holowka, #3) CG components in compartment A (B) is to a lesser extent, and the isochores boundary enrichment is not significant (p-value = 0.14 by chi-square test, Figure 3F and SI). We need to note that the identification of compartments for human genome should be performed with care (SI).

### Intra-domain F-P interactions in 3D chromatin structure

In this section, we investigate the organization of forests and prairies in the 3D chromosome and its biological consequences. We analyzed data for 22 human cells and 20 mouse cells (Supplemental Table 1). The diversity of the dataset allows us to investigate the chromatin structure difference in different cell/tissue types and stages, obtaining information concerning early embryonic development, differentiation and senescence.

To describe the local interactions of forests and prairies, we defined and calculated the forest index (see method). For the 22 human cells, (82.6±4.6) % of forests have positive indices, and (91.2±1.2) % of prairies negative ones. For mouse samples these two values are (92.4±7.5) % and (93.3±1.8) %, respectively. Therefore, the vast majority of chromatin is surrounded by sequences of the same type, indicating that individual forests and prairies separately segregate in space.

However, the extent of this segregation is cell-type specific, independent of the data scale (Figure S15). During early embryonic development, the magnitude of the forest index slightly increases from the early two-cell to the eight-cell stage (Figure 4A), consistent with the re-establishment of TADs and high-order chromatin structures in early development^11, 12, 37, 38^. In differentiation, local F-P interactions tend to increase for both mouse and human. For mouse, the absolute values of average forest indices for both domains decrease from iPSCs to differentiated cells (Figure 4B), especially for more highly differentiated pre-B and macrophage cells. Human cells show similar trend in differentiation. The pluripotent cells have the largest absolute values of forest indices (Figure 4C), while in tissue samples except for liver and spleen which are actively proliferating, the corresponding values are significantly lower (both with p-value = 6.5×10^−4^ by Mann-Whitney U test). Absolute values for forest indices of somatic tissues are uniformly smaller than those of h1-derived pluripotent cells, though in a lineage-specific way (Figure 4D). Even the somatic tissues with the least local F-P interactions, left and right ventricles, still have a weaker segregation than their corresponding h1-derived mesendoderm cells. These results indicate that an increase of F-P interaction accompanying differentiation also occurs in human. Forests and prairies form stronger intra-domain interactions in cell lines and actively proliferating cells (liver and spleen) than normal somatic tissues, though to a lesser extent than pluripotent cells (Figure 4D). The absolute values of average forest indices for cell lines and active cells are larger than those for somatic tissues (both with p-value=2.2×10^−3^ by Mann-Whitney U test), except for cancer cell line PC3 which is close to normal tissue samples.

**Figure 4.**
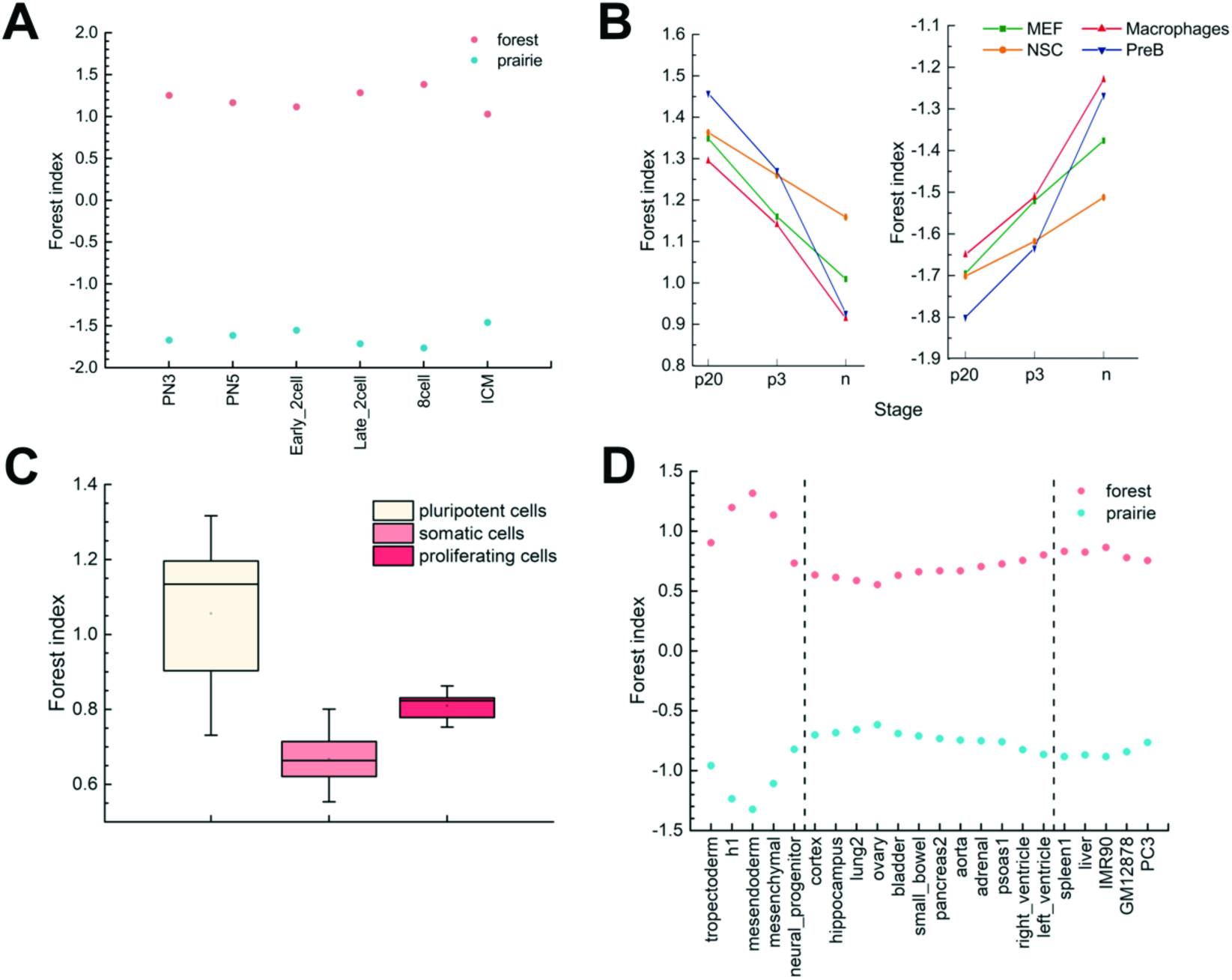
The change of local 3D chromatin structural properties during embryonic development and cell differentiation. (A) Average forest index of forests and prairies in different cells during mouse embryonic development. (B) Average forest index of forests (left) and prairies (right) in different stages during cell differentiation for four mouse cell types. (C) The box plot of forest indices in forest for three types of human cells. (D) Average forest index of forests and prairies in different human samples.

As analyzed above, the gene expression level largely relates with its sequential location on forests or prairies. The spatial packaging of chromatin also regulates gene expression. For 14 human samples with both structural and expression data, genes located in regions with positive forest indices have significantly higher expression levels than those with negative indices (average logarithm FPKM expression level of 0.917 and 0.381 respectively, p-value< 10^−100^ by Welch’s unequal variance t-test). Specifically, even prairies with positive indices (reversed prairies, with an average logarithm expression level of 0.578) are modestly yet significantly more highly expressed than forests with negative indices (reversed forests, 0.522, p-value< 10^−15^ by Welch’s unequal variance t-test). The available mouse samples yielded similar results (SI). Therefore, local spatial environments play an essential role in regulating gene transcription inside these domains.

The reversed forests and reversed prairies of all 22 human samples (defined as merged reversed regions) constitute16.5% and 10.8% of the whole genome, respectively, indicating that forest indices in over 70% of the chromosome are conserved. The merged reversed regions are enriched in F-P boundaries (Figure S16). Noticeably, merged reversed prairies are significantly more enriched in high GC content regions (GC content > 41%, corresponding to H isochores^23^, see SI) compared to all prairies (46.2% versus 18.5%, p-value < 10^−100^ by chi-square test), while merged reversed forests are more enriched in low GC content than all forests (55.2% versus 34.3%, p-value < 10^−100^ by chi-square test). As immune and inflammatory response genes are enriched in forests of low GC content and prairies of high GC content (defined as bivalent regions, see SI and Figure S17), the bivalent sequences may yield a high structure flexibility and possibly a quick response to the environment. Since the mouse genome has longer average domain lengths than the human genome, forests and prairies in mouse are more likely to form intensive intra-domain interactions, resulting in larger absolute values of forest indices and a lower proportion of reversed regions (SI).

Since F-P interactions appear to contribute to chromatin structure re-arrangement and thus dynamic gene regulation, they provide a possible mechanism for cell-specific gene regulation. We then analyzed the function of genes located in cell-specific reversed prairies (reversed prairies for a sample that are shared by less than half of all samples), which are expected to be activated specifically in the corresponding cell type. For example, 27 out of 233 such genes in cortex are related to known brain functions or diseases, as well as tumor suppressors, including ADAM12, which is involved in neurogenesis, DOCK3, which is specifically expressed in the central nervous system (CNS), and HTR7, which relates to various cognitive and behavioral functions (Supplemental Table 2). Functional analysis for cell-specific prairies of h1, GM12878, and IMR90 cell lines also gave cell-type specific results (SI). These results show that the plasticity in forest and prairie interaction is important for the activation of the latter, which correlates strongly with cell differentiation and cell identity establishment.

### 3D chromatin structure features of forests and prairies at the domain level

The forest index is largely dominated by the intra-domain (forest or prairie) interactions and the interactions between nearest neighbors, as a result of the fast decay of the contact probability along the genome. To examine how different domains interact with each other at a longer distance, we calculated the ratio of intra-domain contacts and the inter-domain contacts between domains of the same and different types for forests and prairies, respectively, regarding each domain as one unit (Figures 5A-D). The intra-domain contacts positively correlate to their absolute forest indices across cell-types (with correlations of 0.99 and 0.98 for forests and prairies, respectively), validating that they both reflect the local structural environment. The variations of the intra-domain contacts with cell types are essentially the same for all chromosomes (Figure S18), with an average Pearson’s correlation of 0.976 between every pair of chromosomes. Therefore, we use chromosome 1 as an example in the following analysis.

To quantify the relative inter-domain interactions between the same types relative to that between different types, we define a segregation ratio *R*_*s*_ (see Methods) as the inter-domain contact ratio between the same types and different types. A higher *R*_*s*_ for a sample indicates a stronger segregation of genome domains of the same type. To investigate the different contribution of contacts along genomic distances, we also calculated the contact probability *P*_*c*_(*s*) between domains of the same and different types as a function of genomic distance *s*.

The forest and prairie intra-domain contacts are highly enriched in early embryonic and pluripotent cells for both human and mouse. In early embryonic development from 2-cell stage, intra-domain contacts tend to decrease, coupled with the increase of inter-domain contacts, indicating the establishment of long-range interactions (Figure 5A). The segregation ratio *R*_*s*_ uniformly increases during the development of early embryonic cells except for the early 2-cell stage (Figure 5E), F-P contact probability *P*_*c*_(*s*) also noticeably lowers compared to that of F-F and P-P in this process (Figure S19A). Such changes suggest the increased segregation of the forest and prairie domains from each other, as can be seen from their clustering in the 3D structures reconstructed via Hi-C data (Figure S19A).

**Figure 5.**
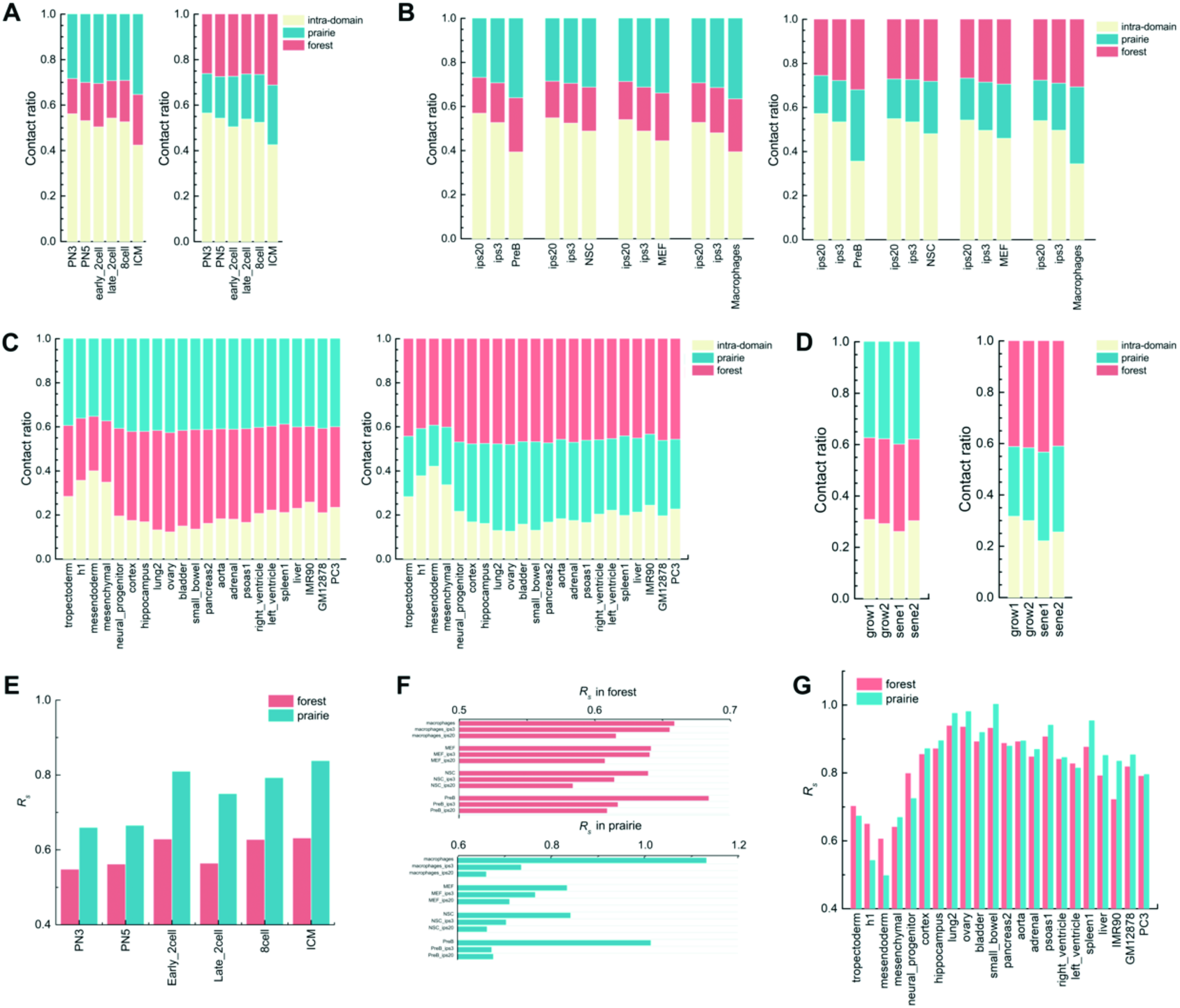
3D chromatin structure features of forests and prairies at domain level. (A-D) The proportions of intra-domain contact and inter-domain contact of the same and different type in forest (left) and prairie (right) (A) for different cells in mouse early embryonic development; (B) for different differentiation stages of four mouse cell types; (C) for different human samples, and (D) for growing and senescent cells. (E-G) The segregation ratio *R*_*s*_ between inter-domain contacts of the same type and different type (E) for different cells in mouse embryonic development; (F) for different differentiation stages of four mouse cell types; (G) for different human samples.

During cell differentiation, forest and prairie domains establish more long-range contacts with the sacrifice of intra-domain ones. In contrast to their corresponding pluripotent cells, decreased intra-domain contacts and increased inter-domain interactions between both the same and different domain types, especially P-P interactions, are clearly observed for human somatic tissues and mouse differentiated samples (Figures 5B, 5C, S19B and S19C). The increase of *R*_*s*_ is also observed in a lineage and cell-type specific way in differentiation (Figures 5F and 5G). For example, the intra-domain contact and *R*_*s*_ of neuronal progenitor are between those for h1 cell line and its corresponding differentiated tissues, cortex and hippocampus (p-value = 0.0025 and 0.002 by Student’s t-test, respectively). The *R*_*s*_ of the highly differentiated mouse pre-B and macrophage cells increases more significantly than other mouse samples. These results support that domains of the same genomic type generally tend to cluster and segregate with cell differentiation, while the interactions between domains of different types increase in a cell-type specific manner.

In proliferating human tissues (spleen and liver) and cell lines, both intra-domain and P-P inter-domain interactions are higher than other somatic tissues (Figures 5C and S19C), consistent with the segregation of prairies. Their reconstructed 3D chromatin structures also exhibit strong domain segregation, in contrast to the structures with highly intermingled forests and prairies of normal somatic tissues (Figure S19C).

We next examined the chromatin structure properties during senescence. Although forest contacts differ relatively little compared to the growing cells, their prairies lose intra-domain contacts and gain P-P inter-domain contacts (Figures 5D and S19D). Specifically, the *R*_*s*_ for all prairie domains in senescent cells are almost uniformly higher than those in growing cells, while the *R*_*s*_ distribution of forests becomes broader with little change of the average value (Figure S20). These results are consistent with previous observations of clustering of H3K9me3 enriched regions and loss of local contacts in senescent cells^15, 39^, and indicate a senescent mechanism involving prairie clustering and segregation from forests.

### Partition of forests and prairies in compartments A and B is related to gene regulation

As discussed above, forests and prairies preferentially locate in compartments A and B, respectively. In addition, we found that during cell differentiation, the proportion of chromosomes belonging to compartment B generally increases except for the NSC cell (Figure 6A), consistent with the growth of the heterochromatin ^40, 41^ and the increase of repressive histone mark covered regions during differentiation ^42, 43^.

**Figure 6.**
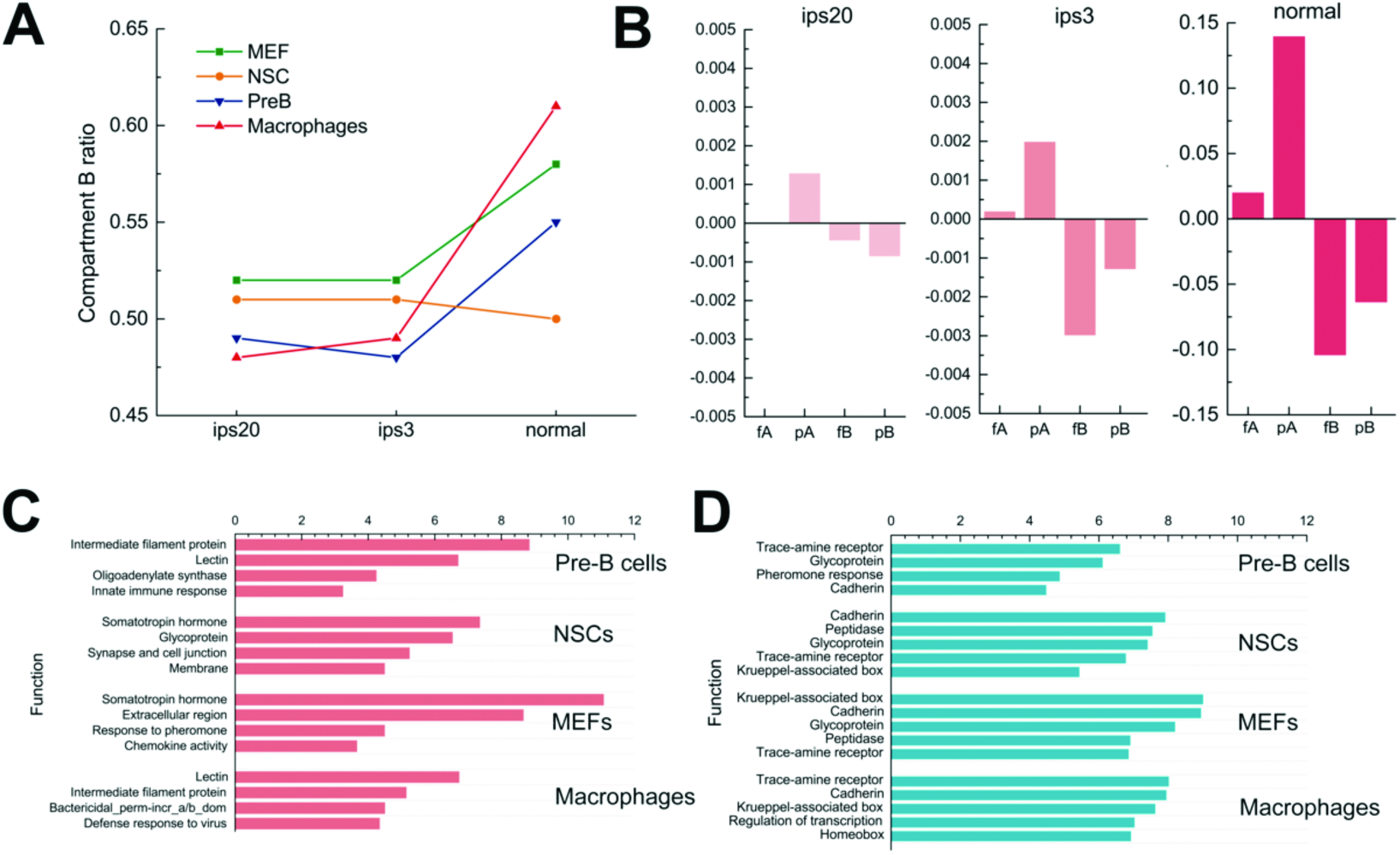
The regulation of cell differentiation viewed from the compartment aspect. (A) The length ratio of compartment B in different differentiation stages for four mouse cell types. (B) Relative gene expression in forests and prairies of compartment A and B, respectively, in different differentiation stages. The relative gene expression level was calculated as the average gene expression level in each component subtracted by the overall average expression level among four samples at the same stage. (C) Functional clustering of genes in prairies of compartment A and (D) forests of compartment B.

To understand how forests and prairies’ positioning in compartments affect the expression of related genes, we divided the genome into fA, fB, pA, and pB components according to the overlap of forests and prairies with compartments A and B. For these four components, genes in pA and fB have the highest and the lowest relative gene expression levels, respectively, regardless of cell types. Their difference increases as cells differentiate (Figure 6B and S21), leading us to speculate that genes in pA and/or fB might possess cell-specific functions. We then clustered gene functions in pA and (Figures 6C and 6D). Genes in pA are related to cell-specific functions (Table S5). For example, pA genes in immune-related macrophages and pre-B cells are enriched in functions such as defense against invading microorganisms, bacterial killing, defense response to virus, and immunity. The main functions of pA genes in MEFs include structural components and immune response to tissue injury. Genes in pA of NSCs correlate well with growth and neuron activities. In contrast, genes in fB are of similar functions for all cell types analyzed, including trace-amine receptor, cadherin, transcriptional repressor domain Krueppel-associated box, homeobox, and peptidase. In summary, the function of genes of prairies located in compartment A are largely cell-type specific, whereas forest genes in compartment B lack cell-type specificity but mainly function in transcriptional repression.

## Discussion

In the present study, we integrated various sources of genetic, epigenetic and 3D structural information to investigate the sequence dependence and cell-type specificity in the formation of 3D chromatin structure. Both human and mouse genomes were found to contain long segments of distinct sequence properties, echoing earlier studies^44^. Based on the distribution of CGIs, we divide the human and mouse genomes into alternative forests (high CGI density) and prairies (low CGI density) with average lengths of megabases. The genome can thus be regarded as an A-B copolymer and the chromatin structure its assembly. Compared to studies using GC content which lead to isochores^23, 24^, the current classification divides the genome into domains of more distinctly different genetic, epigenetic, and 3D chromatin structural properties. Forests enrich not only genes, especially HKGs, but also active histone marks including H3K4me1, H3K4me3 and H3K36me3. Open seas in forests are generally more highly methylated than those of prairies. Prairies in contrast enrich repressive histone marks H3K9me3 and are transcriptionally inactive. The close correspondence of these domains to both TAD and compartments reveals their important roles in 3D chromatin structure formation.

### Chromatin structure change in differentiation and senescence

We found that cell development, differentiation and senescence in human and mouse manifest two types of DNA interaction changes: the F-P interactions which are associated with cell identity establishment and tissue function specification, and the F-F and P-P interactions which lead to domain segregation. The latter strengthens during differentiation and senescence. These two effects influence the chromatin structure profoundly in multiple scales: from intra-domain local contacts related to TAD formation to long-range contacts related to compartments.

In differentiation, F-P interaction coincides with tissue-specific gene functions (Figure 7B), characterized by cell type-specific formation of locally reversed prairie regions and specific prairie penetration into compartment A. In senescence, a subset of forests continues to increase in interactions with prairies. It has been found that HKGs are not involved in interactions with distal regulatory elements^45, 46^, but lineage-specific genes are more dependent on long-range interactions and tend to form complex regulatory networks^45, 47^, thus sensitive to spatial environments. Since transcription factors (TFs) such as YY1 are heavily involved in the establishment of the gene regulation network^48^, we strongly suspect that these F-P interactions are mediated by the binding of proteins (TFs) to the interacting forests and prairies, and call for studies in this direction.

**Figure 7.**
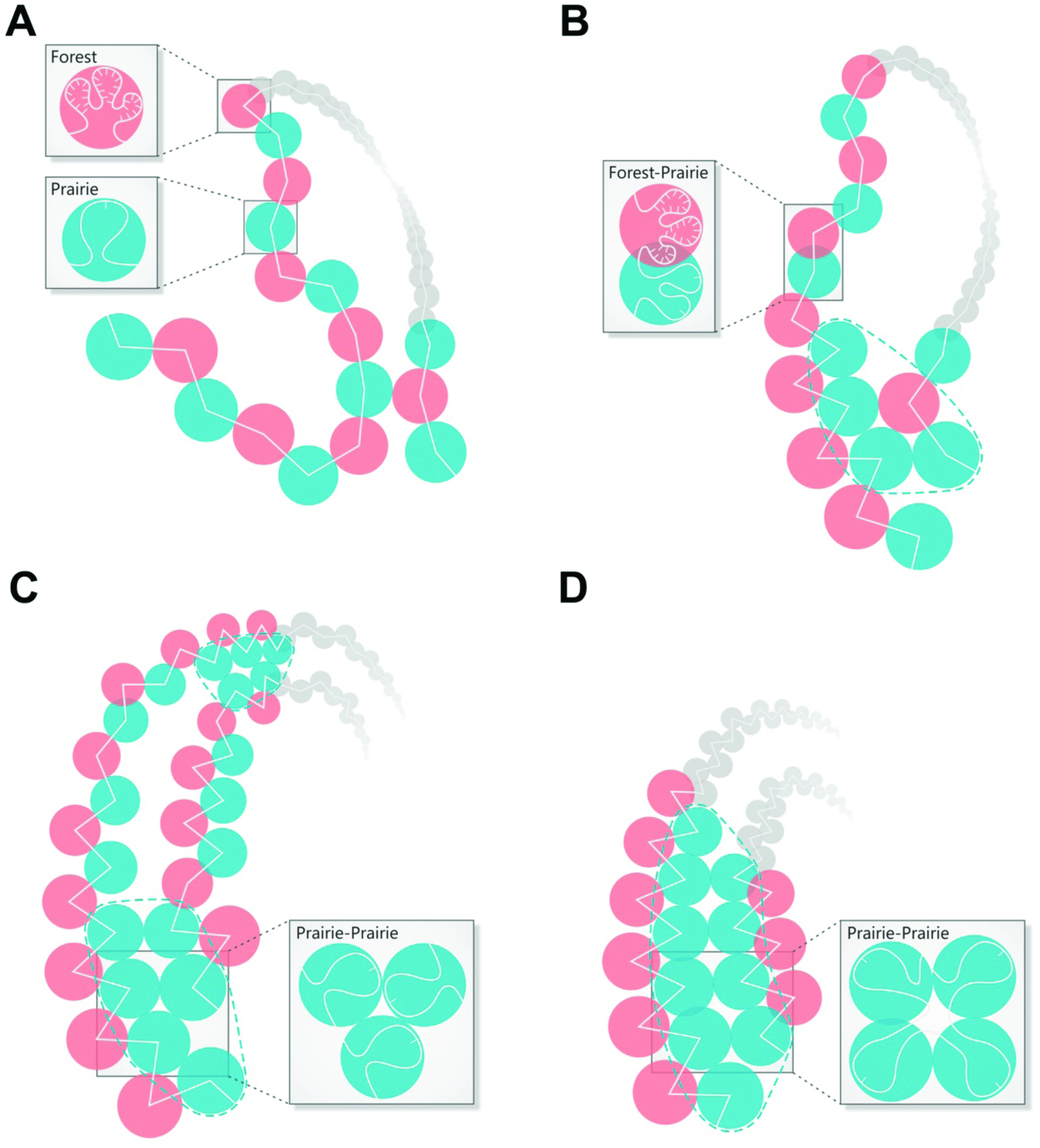
A schematic picture of the forest-prairie separation in (A) pluripotent cells, (B) somatic cells, (C) senescent cells, and (D) actively proliferating cells.

On the other hand, due to their different sequence and possibly physical properties as discussed below, forests and prairies have a natural tendency to segregate from each other. In early embryonic development, intra-and inter-domain segregation are observed to progress from a relatively homogeneous state (Figure 7A), along with the TAD establishment and compartment formation^11, 12^. In differentiation, domains of the same types cluster and heterochromatin accumulates (Figure 7B). These trends continue in senescence, in which the segregation becomes dominant over F-P intermingling (Figure 7C). Consistent with our observations, Chandra et al^15^ found that senescence is characterized by the spatial clustering of constitutive heterochromatin and H3K9me3 repressive histone marks, as well as the loss of intra-domain interactions of genomic segments of low GC content. We also note here that in mitosis, the position of a given chromosome territory (CT) reshuffles and neighboring CTs can vary between mother and daughter cells.^49, 50^ However, despite this cell-cell variation, regions with similar transcriptional activity tend to interact more frequently with each other, which extends beyond a single chromosome, both for active loci and inactive regions^1, 51^.

Similar to embryonic and pluripotent cells, proliferating tissues and immortalized cell lines also possess segregated chromatin structures (Figure 7D). However, different from normal somatic tissues, the P-P contact probability in proliferating tissues and cell lines is significantly higher than that of F-F, suggesting global segregation of prairie domains. Notably, despite that cell lines and senescent cells both have more frequent P-P contacts than those of F-F, cell lines are more enriched in intra-domain contacts for both forests and prairies, and senescent cells are short of intra-prairie contacts. Chromosomes of cell lines are more segregated at both intra-domain and inter-domain levels, but those of senescent cells only exhibit elevated inter-domain interactions. The domain segregation mechanisms are thus different for proliferation and senescence.

The chromatin structure change in differentiation and senescence resembles a “phase separation” mechanism that is characterized by the cell-specific removal of prairies from the active chromatin domains. The domain segregation contributes to stable differentiation, and possibly irreversible aging at the same time. This chromatin stability is consistent with the need of ATP associated chromatin remodeling factors in reprogramming^52^^-^^54^. Meanwhile, life might have evolved to cope with the irreversible chromatin structure segregation and the corresponding aging by introducing fertilization which yields globally F-P mixed and homogeneous chromatins. This study, along with earlier analyses^11, 12^, showed that the early embryonic chromatins are characterized by weak domain segregation, small heterochromatin, CpG demethylation, and less repressive histone marks. Whereas differentiation and senescence are associated with the non-specific segregation of forests and prairies into different compartments, as well as the establishment of TF-mediated specific genomic interactions^48, 55^.

### Correlation between epigenetic marks and F-P “phase separation”

The F-P open sea methylation difference in different cells corresponds to their different degrees of segregation. The methylation difference is more prominent for immortalized cell lines, especially for IMR90, which shows a stronger forest-prairie separation, than that for normal somatic cells. Meanwhile, pluripotent cells with less segregation also have a smaller methylation difference between forests and prairies. Interestingly, cancer cells show large F-P methylation differences. It is thus reasonable to speculate that cancer chromatin associates with a large F-P separation. The one cancer cell line data available to us does show more apparent phase separation than typical somatic tissues, but similar to the liver sample. Incidentally, the liver cell also shows an F-P methylation level difference larger than other somatic cells and a power law scaling of the methylation correlation which is in between somatic and cancer/immortalized cells (−0.26±0.02 for normal somatic cells, −0.06±0.02 for cancer samples, and −0.083 for liver, respectively)^32^. We hope that more systematic Hi-C studies will shed light onto the chromatin structure change in oncogenesis.

We also want to point out that the megabase-scale forests and prairies are enriched with active histone marks H3K4me3, and repressive H3K9me3 mark, respectively. However, another repressive mark H3K27me3 was found to show more cell-specificity and is relatively more enriched in forests, consistent with them being localized inside compartment A^56^ and their co-localization with forest-enriched H3K4me3 to form bivalent regions^57, 58^. Since the establishment of H3K4me3 and H3K27me3 marks, and the local aggregation of H3K27me3 decorated and polycomb complex occupied domains are important for differentiation^56, 59^, a detailed analysis of the association and segregation of genome domains marked by H3K4me3, H3K27me3, and H3K9me3, as well as their roles in differentiation, is of great interest.

### The possible physical mechanism and consequence of “phase separation”

As we have observed, the F-P boundaries overlap well with the TAD boundaries. The former depends only on the DNA sequence and the latter is largely conserved among different cell-types ^4^. Although structural proteins are important for the formation of TADs and the CTCF binding sites can be used to predict Hi-C contact maps^60, 61^, the loss of cohesin did not lead to the disruption of 3D chromatin structure^19^, and a quarter of TAD boundaries show no evidence of CTCF binding^4^. It is thus possible that more fundamental causes exist for TAD formation. Recent study on senescent cells suggests that epigenomic remodeling and chromatin structure changes could be discrete events^39^. The breakdown and re-establishment of TADs as well as the reshuffling of chromosome territory during mitosis in which factors like HP1 and polycomb proteins are excluded from the sequence^49, 50, 62, 63^ also suggest a more intrinsic mechanism for chromatin structure sustainability than protein binding. These observations all promote us to propose that the segmented DNA sequence provides a more fundamental driving force for the formation of chromatin structure. The sequentially distinct forests and prairies have the natural tendency to form largely self-interacting, thus insulated local structures of different contact patterns, which further leads to two major compartments in nuclear, the more open compartment A, and the less accessible compartment B.

It then becomes intriguing to speculate on the origin and biological consequences of the physical forces that might lead to the segregation of prairies from forests during development and aging. As prairies are of low DNase signals and are highly occupied by histones, which neutralize the charges and make the genome less polar, the prairies thus are expected to be less soluble in an aqueous environment. It was recently observed that HP1a protein which is characteristic of heterochromatin undergoes liquid-liquid demixing in vitro, and that heterochromatin exhibits liquid-phase separated dynamics^64^, consistent with the segregation of prairie sequences from forests being physically driven by a phase separation mechanism. It is known that the solubility of hydrophobic particles decreases with temperature. Therefore, an increase of temperature could induce the “precipitation” of the prairies, whereas forests are more likely to remain in the aqueous phase. Cell differentiation is thus accompanied by the growth of the less polar and less active phase, which is closely related with compartment B and heterochromatin. The growth of this phase is consistent with the observation that the chromatin of pluripotent cells tends to be relatively homogeneous with less heterochromatin which becomes more prevalent in differentiated cells ^40, 41^, as well as the spread of repressive H3K9me3 and H3K27me3 histone marks in differentiation^42, 43, 65^. The phase-segregation is enhanced as cells proliferate, heading to a more stable structure. Tissue (or cell)-specific domains of the prairies remained in the open chromatin through mediators like tissue-specific transcription factors contribute to cell identity establishment.

As temperature may affect the domain segregation, one would also expect the chromatin structure to vary with tissues of different temperatures. Liver is an organ of the highest temperature^66^, thus is expected to exhibit a strong F-P segregation. Indeed, compared to other somatic tissue samples, the Hi-C data of liver is characterized by stronger intra-domain interactions and distant PP interactions. Its reconstructed 3D chromatin structure also shows a high degree of domain segregation. Meanwhile, the liver sample also shows a large F-P methylation difference and its scaling power of methylation level correlation function is accordingly much less negative than other somatic cells. In contrast, the brain chromatin sample is characterized by strong F-P local interactions and weak domain segregation, with an open reconstructed chromatin structure (Figure S20). Consistently, the scaling power in the methylation correlation function of brain samples is more negative than normal somatic tissues, and its F-P methylation difference is small. In addition, the scaling power of brain tissues becomes more negative in neurodegenerative diseases, indicating a possible association of domain segregation and temperature effects to these diseases (SI).

### Implications on species differences

As discussed earlier, not all genomes are of the same mosaic property. In fact, CpG deficiency appears to emerge only at a certain stage of evolution. Based on the postulation that spatial domain segregation stabilizes differentiation, genomes with lower mosaicity (e.g., with more uniform CG distribution, such as bacteria and plant *Arabidopsis*^67, 68^) would have weaker domain segregation and less stable differentiation. This model predicts that it is easier for cells of species with less mosaic DNA properties to reprogram.

The sequential difference between different species may also correlate to their different responses to the environmental temperature. As the phase-separation model suggests, a narrower temperature range is required for proper functioning of species with a stronger mosaic genome. In fact, warm-blooded animals in general have more apparent sequence heterogeneity than coldblooded animals^69^, and correspondingly tend to keep their body temperatures at a species-specific narrow range ^70^. Reptiles, on the other hand, have a much broader body temperature range^71^.

The phase separation argument also indicates that temperature strongly affects the chromatin structure remodeling, thus affects embryonic development and differentiation. In fact, it is known that temperature is an important factor for the embryogenic development. High temperature relates to the abnormal development and low temperature slows down or even terminates the development^72, 73^. Seed germination also requires appropriate temperatures^74, 75^. Among many other examples, the incubation temperatures of ovipara are closely related to the state of their offsprings^76^, and the sex of alligator can be affected by the incubation temperature^77^.

As another possible effect, a lower body temperature is expected to slow down the senescence of the cells as domain segregation is related to senescence. Low temperatures have been found to relate to the expansion of the life span of rotifera, mice models and human^78^^-^^80^. Chromatin segregation is also seen in the immortalization and possibly also oncogenesis of cells. Interestingly, cancers are also less likely found in animals of lower or variable body temperatures, such as elephant and naked mole rat, than those with high or steady body temperatures^81^^-^^84^. Such correlations between temperature, chromatin segregation, and diseases, are far from being understood and worth careful investigation.

In summary, important correlations have been identified between human and mouse DNA sequences, their 3D chromatin structures, epigenetics and various biological functions. Chromatin structure changes involving development, differentiation and disease were explained based on a rather simple theoretical framework which involves the genomic mosaicity and the segregation of the genomic domains of different sequential and physical properties. It is intriguing to examine in a more systematic way the roles of genomic information and phase separation in the chromatin structure formation for different species, at different cell and disease states, and under different environmental conditions, especially at different temperatures.

## Supplemental Information

Supplemental Information is available in the online version of the paper

## Acknowledgement

We thank Xiaoliang Sunney Xie and Xing Chen for helpful discussions. The results shown here are partly based upon data generated by the TCGA Research Network: http://cancergenome.nih.gov/. We are grateful to Manel Esteller for providing the methylome of neurodegenerative diseases. This work was supported by National Natural Science Foundation of China [21573006, 21233002 and 91427304 to Yi Qin Gao, U1430237 to Lijiang Yang] and the National Key Research and Development Program of China [2017YFA0204702 to Yi Qin Gao].

## Author Contributions

Yi Qin Gao designed the research. Sirui Liu, Ling Zhang, Hui Quan and Hao Tian performed the research. Sirui Liu, Ling Zhang, Hui Quan and Yi Qin Gao wrote the manuscript. Luming Meng contributed to the analysis of the data, Huajie Feng and Lijiang Yang helped with the manuscript writing.

## Competing Interests

The authors declare that they have no competing interests.

